# Dynamics and functional roles of splicing factor autoregulation

**DOI:** 10.1101/2020.07.22.216887

**Authors:** Fangyuan Ding, Christina Su, Ke-Huan Kuo Chow, Michael B. Elowitz

## Abstract

Non-spliceosomal splicing factors are essential, conserved regulators of alternative splicing. They provide concentration-dependent control of diverse pre-mRNAs. Many splicing factors direct unproductive splicing of their own pre-mRNAs through negative autoregulation. However, the impact of such feedback loops on splicing dynamics at the single cell level remains unclear. We developed a system to dynamically, quantitatively analyze negative autoregulatory splicing by the SF2 splicing factor in response to perturbations in single HEK293 cells. Here, we show that negative autoregulatory splicing provides critical functions for gene regulation, establishing a ceiling of SF2 protein concentration, reducing cell-cell heterogeneity in SF2 levels, and buffering variation in SF2 transcription. Most importantly, it adapts SF2 splicing activity to variations in demand from other pre-mRNA substrates. A minimal mathematical model of autoregulatory splicing explains these experimentally observed features, and provides values for effective biochemical parameters. These results reveal the unique functional roles that splicing negative autoregulation plays in homeostatically regulating transcriptional programs.

## Introduction

More than 90% of a typical eukaryotic genome undergoes alternative splicing, producing multiple mRNA isoforms and expanding proteome diversity (Barbosa-Morais et al., 2012; Merkin et al., 2012; Nilsen and Graveley, 2010; Pan et al., 2008; Wang et al., 2008). Mis-splicing can cause diverse physiological effects and lead to disease (Faustino and Cooper, 2003; Kalsotra and Cooper, 2011; Scotti and Swanson, 2016). Alternative splicing is controlled by many distinct components (Black, 2003; Lee and Rio, 2015), including non-spliceosomal splicing factors (Jangi and Sharp, 2014), the splicing code (Barash et al., 2010; Culler et al., 2010), RNA secondary structures (McManus and Graveley, 2011), RNA polymerase speed (Fong et al., 2014), and epigenetic regulation (Luco et al., 2011).

Of these regulators, non-spliceosomal splicing factors play a unique role by modulating splicing activity in a concentration-dependent manner (Black, 2003; Shin and Manley, 2004; Wang and Burge, 2008). Splicing factors fall into two main, conserved families, serine-arginine rich (SR) proteins and heterogeneous nuclear ribonucleoproteins (hnRNPs), and are found across diverse tissue types and species (Dreyfuss et al., 1993; Manley et al., 1996; Zahler et al., 1992), ranging from *Schizosaccharomyces pombe* (Shepard and Hertel, 2009) to *Arabidopsis* (Kalyna et al., 2006). SR or hnRNP proteins modulate alternative splicing of large and diverse sets of target genes (Long and Caceres, 2009; Wang and Manley, 1995; Zhou and Fu, 2013), and are implicated in diverse disease processes (Anczuków and Krainer, 2016; Geuens et al., 2016). Maintaining splicing factor homeostasis is thus critical for cellular function.

The control of splicing factor level is commonly achieved through negative autoregulatory splicing (Kalsotra and Cooper, 2011; Lareau et al., 2007; Ni et al., 2007). Specifically, splicing factors alternatively splice their own pre-mRNA to unproductive isoforms, either containing premature termination codons (Wollerton et al., 2004), or introducing new junctions in the 3’ untranslated region (Sureau et al., 2001), to trigger degradation by RNA surveillance pathways (Maquat, 2004). Overexpression of splicing factors promotes unproductive splicing (Sureau et al., 2001), leading to negative autoregulation. Previous work investigated many aspects of negative autoregulation, including associated highly or ultra-conserved sequence motifs (Lareau et al., 2007; Ni et al., 2007) and related nonsense-mediated mRNA decay triggered by this regulatory mode (Hug et al., 2016; Ni et al., 2007). Nevertheless, the basic question of how autoregulation plays out dynamically at the single cell level remains unclear.

In other contexts, negative autoregulatory transcriptional feedback is known to speed response times and promote robustness to perturbation (Becskei and Serrano, 2000; Elowitz et al., 2002; Rosenfeld et al., 2002). However, negative splicing regulation potentially has unique features compared to transcriptional feedback. For instance, rather than operating at a fixed number of DNA binding sites, splicing factors can operate at diverse ‘loads’ of pre-mRNA substrates from their own and other target genes in the cell (Figure 1). Due to the effects of stochastic gene expression (Raj and van Oudenaarden, 2008), total substrate amounts can vary between cell states or over time. This provokes the question of what role splicing autoregulation might play in enabling homeostatic control of splicing factor levels and accelerating responsiveness to changes in substrate.

**Figure 1:**
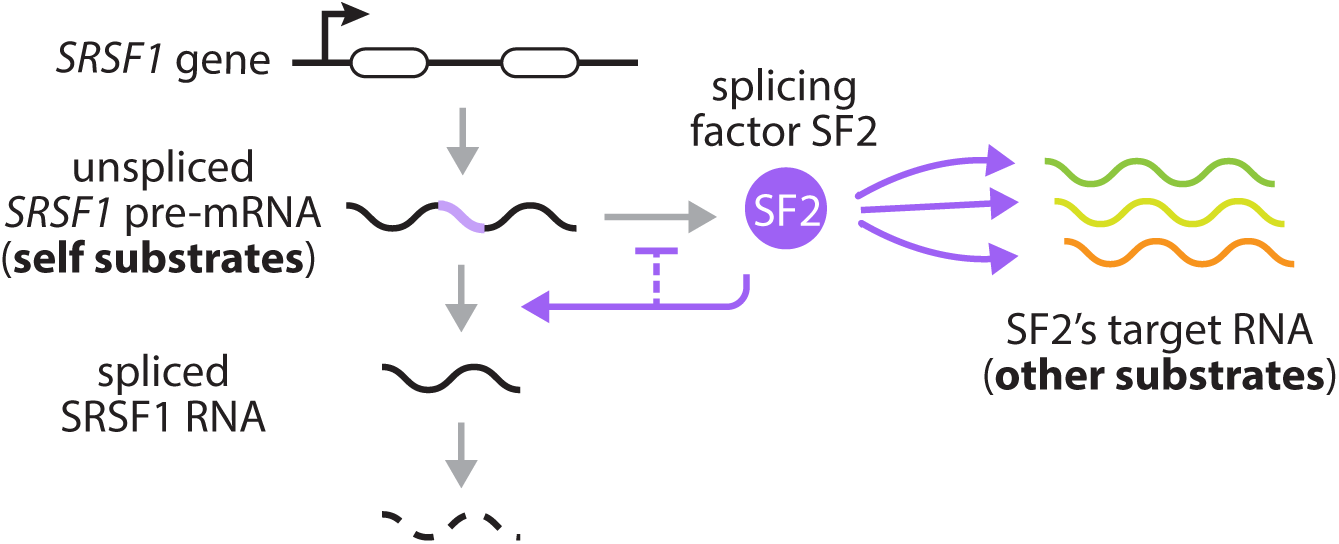
Splicing factors negatively autoregulate their own synthesis by promoting unproductive splicing of their own transcripts. They also operate under variable ‘loads’ of substrate pre-mRNA produced by other genes (on the right). Each splice factor pre-mRNA molecule can be spliced to remove introns (without light purple isoforms) or left unspliced (with light purple isoforms). Intron removal can lead to degradation through RNA surveillance pathways, while transcripts with retained introns are translated to produce more splicing factor (dark purple). Feedback occurs when splicing factors enhance intron removal from their own pre-mRNA, thus negatively regulating their own expression. Apart from their own transcripts, splicing factors additionally act on transcripts produced by other genes. The relative abundance of substrate pre-mRNAs can affect the allocation of splicing factors among transcripts, thereby influencing the dynamics of splicing negative autoregulation.

To address these questions, we developed a system to dynamically and quantitatively investigate splicing negative autoregulation at the single cell level. We focused specifically on SF2, the protein product of the *Serine/arginine-rich splicing factor 1* (*SRSF1*) gene, which is widely expressed in distinct cell types and regulates the splicing pattern of many important genes (Anczuków et al., 2012; Karni et al., 2007; Li and Manley, 2005; Sanford et al., 2009). By tracking SF2 accumulation using time-lapse movies with a machine learning based image analysis system, and analyzing SF2 levels together with flow cytometry and RT-qPCR/PCR, we found that negative autoregulatory splicing can buffer SF2 concentration, achieve ∼50% less cell-cell heterogeneity in expression and response rate upon perturbation of its own pre-mRNA levels, and enable adaptation to total substrate ‘load’ at both a single cell level and across 53 human tissue types. We also demonstrated how negative splicing autoregulation can maintain its own dynamics and how it adapts to perturbation in a minimal model. Together, these results quantitatively explain the single cell dynamics of negative splicing autoregulation and reveal its functional role and impact as a regulatory circuit.

## Results

### SF2 maintains homeostasis through negative splicing feedback

We set out to investigate negative autoregulatory splicing by engineering two HEK293 cell lines expressing the *SRSF1* gene with or without autoregulation. We site-specifically and stably integrated *SRSF1* as a single ectopic copy, using either genomic sequence (gDNA, autoregulated) or cDNA sequence (unregulated), under the control of a doxycycline(dox)-inducible CMV promoter (Figure 2A Top). Because SF2 is essential for cellular physiology (Li and Manley, 2005; Wang and Manley, 1995), the endogenous copy of *SRSF1* was left intact in both cell lines. To distinguish ectopic and endogenous SF2, we fused a fluorescent protein, Citrine, at the 5’ end of the ectopic copies (Figure 2A Top). Addition of Citrine did not affect SF2 RNA or protein levels (Supplementary Figures 1A,B). Importantly, the splicing pattern of Citrine-fused *SRSF1* remained similar to that of the endogenous copy, with 5 isoforms (Supplementary Figure 1A), including the productively translated functional isoform 1 as well as the unproductive isoforms 2-4 (Sun et al., 2010). Like endogenous SF2, ectopic Citrine-fused SF2 can down-regulate total SF2 expression by promoting unproductive splicing to isoforms 2-4 (Supplementary Figure 1B).

**Figure 2:**
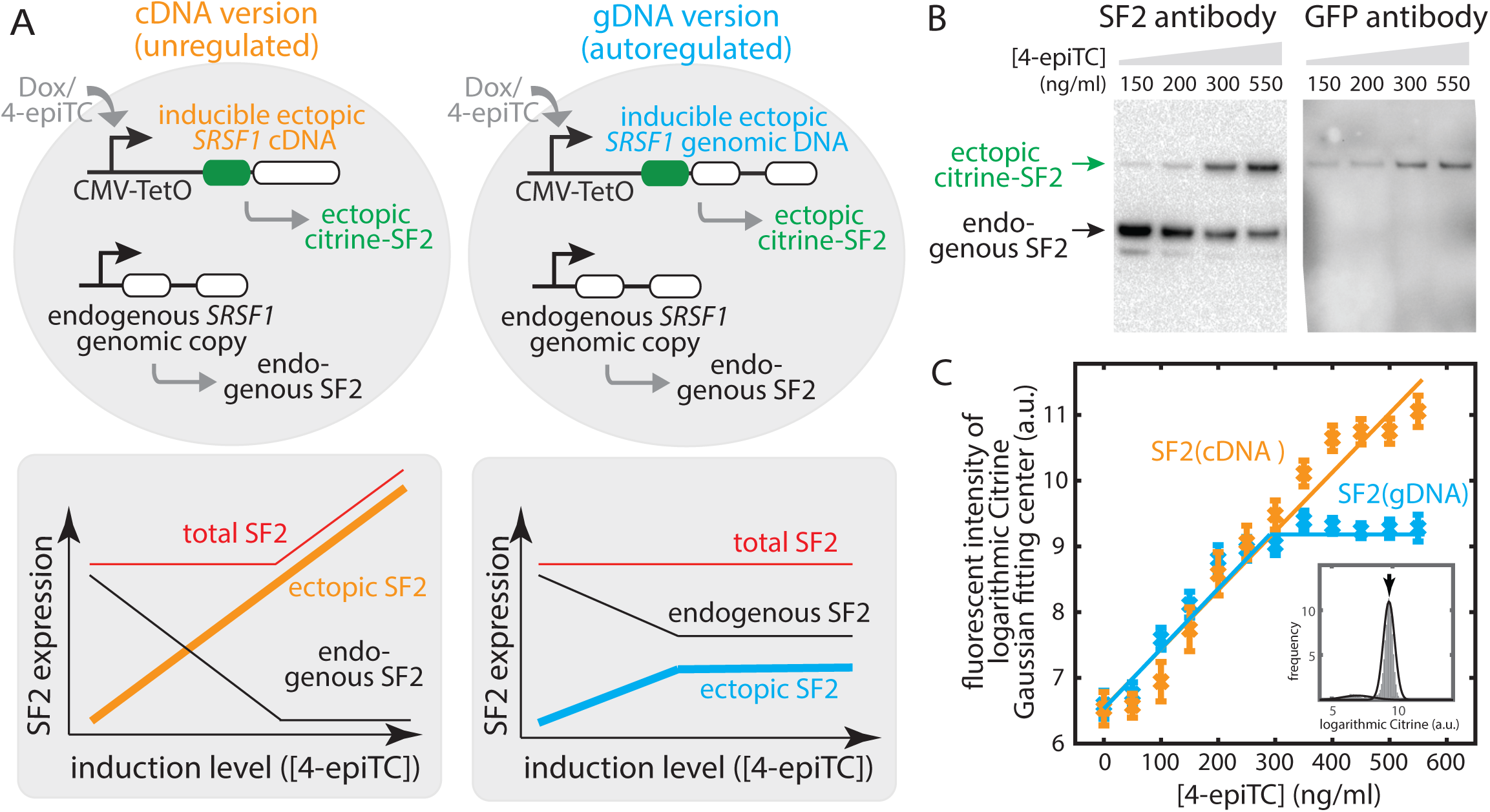
Negative splicing autoregulation establishes a ‘ceiling’ for SF2 protein concentration in response to its own pre-mRNA substrate perturbations. (**A**) (Top) We designed two cell lines: one transfected with a Citrine fused SRSF1 cDNA (i.e. unregulated, with no intron, shown in orange), the other transfected with a Citrine-fused genomic SRSF1 DNA (i.e. autoregulated, shown in blue), both under an inducible Tet-On CMV promoter and stably integrated into the fixed locus of Flp-In T-REx HEK293 cell lines. (Bottom) Expected outcomes (schematic): For the cDNA version, increasing ectopic SF2 protein level should down-regulate endogenous SF2 production via splicing feedback (black curve). When the endogenous copy saturates its ability to buffer SF2 overexpression, the total SF2 level overshoots (red curve). By contrast, for the gDNA cells, due to the negative splicing autoregulation of both the ectopic copy (blue curve) and endogenous copy (black curve), the total SF2 level should remain constant (red curve), across a broader range of induction levels. (**B**) Western blot shows that the endogenous SF2 level decreases with the increasing expression of the ectopic copy. We induced cDNA cells at different 4-epiTC concentration for 24hrs (see gDNA version Western blot in Supplementary Figure 2A). Anti-SF2 antibody (ab133689) staining shows two bands: the top band indicates the ectopic copy with fused Citrine (verified by staining Citrine using anti-GFP monoclonal antibody (right)), the bottom band indicates the endogenous copy. (**C**) Flow cytometry data shows that ectopic SF2 reaches a ceiling (blue curve) with negative splicing autoregulation (i.e. gDNA version), but not with the cDNA version. The two cell lines (A) were induced at different 4-epiTC concentration for >24hrs and analyzed by flow cytometry. Mean expression levels were extracted from Gaussian fits (Supplementary Figure 3) to represent the ectopic SF2 level. Solid lines are guides to the eye to highlight the ‘ceiling’ behavior. Error bars represent standard error of the mean from 9 experimental replicates.

Having established the SF2(gDNA) and SF2(cDNA) cell lines, we next investigated quantitatively how SF2 modulates its own expression. In both cell lines, autoregulatory feedback on the endogenous SF2 gene is expected to maintain constant SF2 protein levels across a modest range of ectopic expression (Fig. 2A, lower panels). However, this buffering effect should saturate in the cDNA cell line once endogenous SF2 is fully depleted (black curve), leading to increased total SF2 levels (orange curve). By contrast, for SF2(gDNA) cells, both SF2 copies (ectopic and endogenous) are autoregulatory. Therefore, the total SF2 expression should remain the same (red curve), with a stable ‘ceiling’ of total ectopic SF2 (blue curve).

Consistent with these expectations, the buffering effect from negative autoregulatory splicing could be observed in both cell lines. Inducing ectopic SF2 with different concentrations of dox or the weaker affinity inducer 4-epiTC (4 epimers tetracycline, an analog of dox) produced a broad range of ectopic SF2 protein expression (Figure 2B,C). Concomitant with the increase of ectopic SF2 levels, we observed a decrease in endogenous SF2 levels in both SF2(cDNA) and SF2(gDNA) cell lines (Western blot in Figure 2B, Supplementary Figure 2A). We also observed a saturation of the buffering effect of endogenous copy in SF2(cDNA) cells: At ∼100ng/ml dox induction, all endogenous pre-mRNAs were spliced to unproductive isoforms 2-4 by 50 hours (Supplementary Figure 1B). These results, which are consistent with previous work done in HeLa cells (Sun et al., 2010), show that the autoregulatory feedback loop is active, and can buffer variations in SF2 expression levels.

To obtain a more quantitative view of this behavior, we used single cell flow cytometry to analyze SF2 protein levels in individual cells (Figure 2C). We induced ectopic SF2 across a range of levels, fit the resulting data to a simple log-normal distribution with background (Supplementary Figure 3), and extracted the mean log expression level for each condition. As expected, ectopic SF2(cDNA) increased monotonically with induction level (orange curve). By contrast, SF2(gDNA) reached a ‘ceiling’ at an induction level of 300ng/ml 4-epiTC (blue curve).

The ‘ceiling’ of expression reached by SF2(gDNA) allows quantification of the transcription strength of the endogenous *SRSF1*. Because the main difference between endogenous *SRSF1* and ectopic SF2(gDNA) is their promoter, different transcriptional strengths can be calibrated by their effects on the ectopic induction level at which the ‘ceiling’ appears. Here, we quantified levels of ectopic and endogenous SF2 via Western blot (Supplementary Figure 2B). Specifically, quantification of gel band intensity, correcting for relative protein size, indicated that endogenous SF2 transcription is ∼3 times as strong as the fully induced CMV promoter (Qin et al., 2010).

Together, these results provide a system in which autoregulated and unregulated SF2 can be compared quantitatively, and show that negative autoregulatory splicing buffers SF2 concentrations at the steady state. We next sought to use this system to study the dynamics of negative autoregulatory splicing in individual cells in response to induction of the ectopic constructs.

### A deep learning system allows label-free single cell movie tracking

Tracking SF2-Citrine in individual cells can be challenging because fluorescence levels are initially low and indistinguishable from background. Therefore, we applied a deep-learning network, BF-Net (Figure 3A), to achieve label free segmentation of cells from brightfield (DIC) images (Christiansen et al., 2018; Szegedy et al., 2015). This method avoids the need to integrate an additional constitutively expressed fluorescent protein into the cell, minimizing cell engineering, and reducing phototoxicity during imaging. However, brightfield images can vary in background, contrast, and evenness of illumination. To address these issues, we trained a single BF-Net (based on GoogLeNet (Szegedy et al., 2015)) by gathering brightfield images from varied imaging conditions as input (left panel in Figure 3A) and corresponding strongly induced fluorescence images of the same cells as ground truth (right panel, Figure 3A).

**Figure 3:**
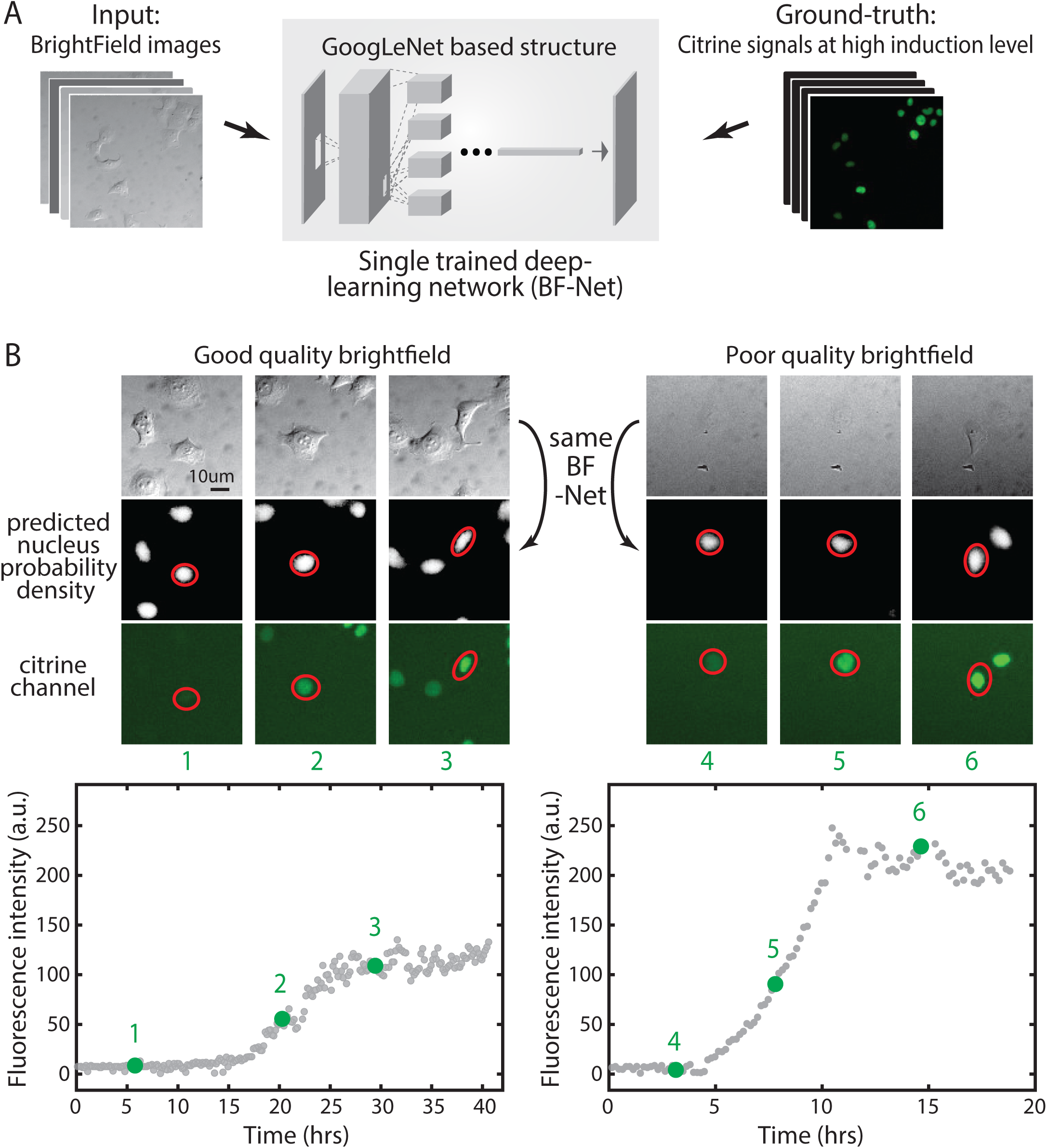
Deep-learning network enables tracking of SF2 accumulation in individual cells over time. (**A**) We trained a single deep-learning network (BF-Net) using ∼150 bright field (DIC) images with their correspondent Citrine signal as the ground truth for learning nuclear location. (**B**) The trained BF-Net predicts nucleus images (middle row) from brightfield images (top row) for two different example time-lapse single-cell traces (left and right panels). Note similarity of predicted fluorescence and actual fluorescence on later images, where visible. Time points are indicated by green numbers (compare with plots). Two movies show diverse brightfield background and contrast, but a trained BF-Net works on both. Red circles represent cell segmentation based on BF-Net predicted nuclear probability. Left and right traces are the SF2(cDNA) cell line induced at t=0 with 200ng/ml 4-epiTC or 100n/ml doxycycline, respectively.

The trained BF-Net was able to directly segment cells in various brightfield images with diverse contrast and illumination conditions. Two examples are shown in Figure 3B. BF-Net predicted nucleus location, based only on brightfield images, correlates strongly with Citrine signal (Figure 3B, position 3 and 6), even under conditions that were difficult to visualize by eye (Figure 3B, position 4-6). We used the predicted nuclear probability density to segment cells and extract fluorescent intensity from the Citrine channel, even for cells with very weak signals (Figure 3B, position 1 and 4). We then applied a previously described cell tracking algorithm to obtain dynamic traces of single cell Citrine fluorescence (Bintu et al., 2016; Singer et al., 2014). Together, this deep-learning-enabled protocol provides a simple and general method for label-free single cell fluorescence tracking.

### Negative autoregulatory splicing reduces cell-cell heterogeneity in SF2 levels and responsiveness

To investigate the dynamics of negative autoregulatory splicing, we compared SF2(gDNA) dynamics to those of unregulated SF2(cDNA), at the single cell level. We induced both cell lines at time 0 under low (200ng/ml 4-epiTC) and high (100ng/ml dox) induction levels, recorded DIC and fluorescence images over time, and reconstructed ∼200 traces of single cell Citrine signals for each cell line under each induction level.

Negative autoregulation at the transcriptional level was previously shown to accelerate response times and to reduce cell-cell heterogeneity (illustrated in Figure 4A) (Alon, 2006; Rosenfeld et al., 2002). Our data show that negative autoregulatory splicing has a similar impact: SF2(gDNA) cells reached steady-state levels faster than SF2(cDNA) cells (Figure 4B).

**Figure 4:**
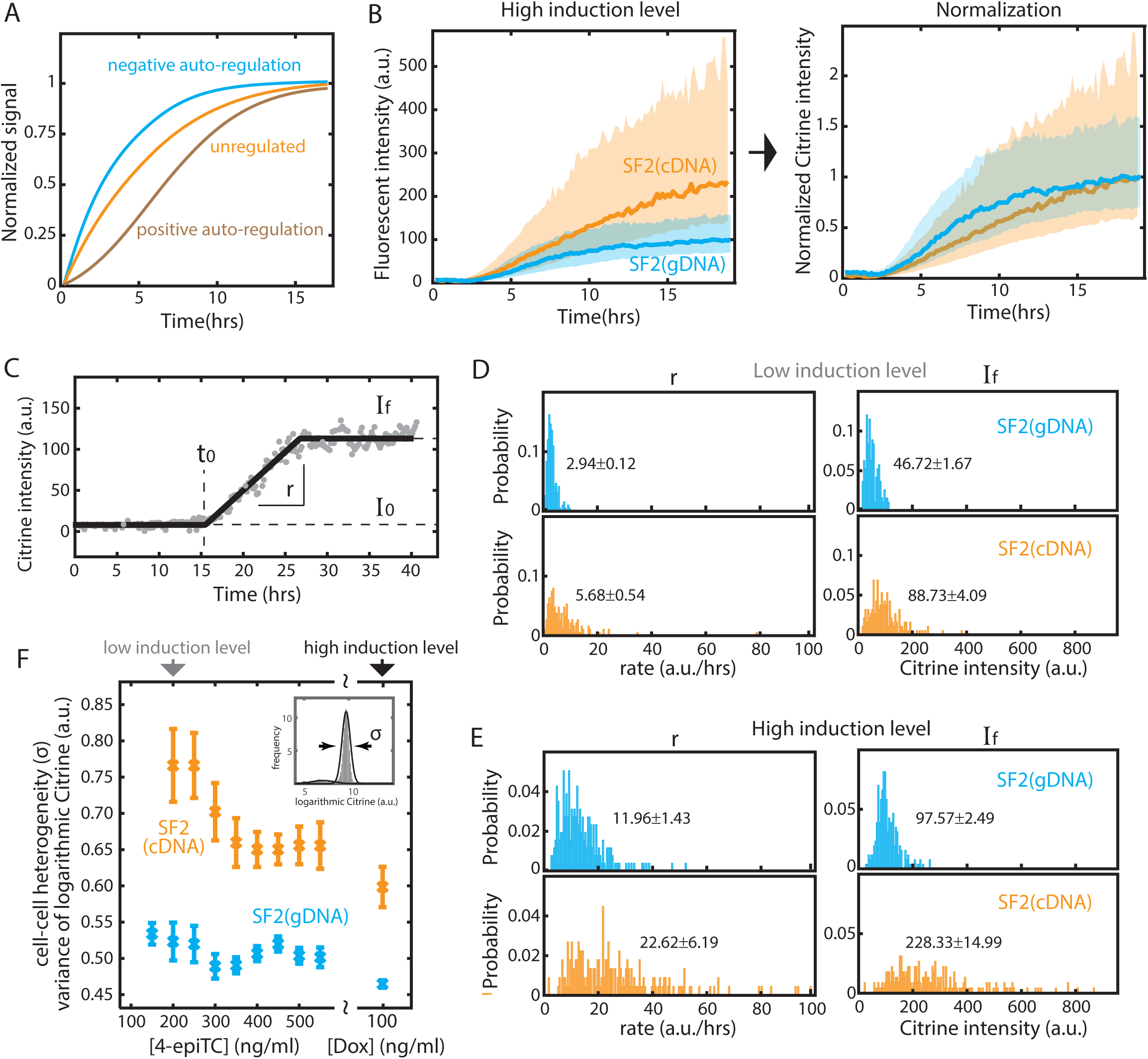
Splicing negative autoregulation reduces cell-cell heterogeneity in both the level and response rate of SF2 protein. (**A**) Negative feedback accelerates response time, in comparison to unregulated, or positive feedback (schematic). (**B**) Negative auto-regulatory splicing speeds response rate and reduces cell-cell heterogeneity at high induction level (100ng/ml dox). (Left) Solid curves are the median of 257 SF2(cDNA) and 223 SF2(gDNA) single cell traces. The shade represents the standard deviation of the mean. (Right): The curves are normalized to final expression. (**C**) We fit four parameters to characterize time-lapse movie traces. I0 represents background Citrine intensity and auto-fluorescence. If represents the final (steady state) Citrine level. t0 denotes the time point when Citrine signal (i.e. ectopic SF2 level) surpasses background. r represents the response speed (slope) from I0 to If. The example movie trace is the same as in Figure 3 (left). (**D**) Distributions of rate and final intensity for 191 gDNA and 188 cDNA traces with 200ng/ml 4-epiTC added at time 0 (See I0 and t0 distribution in Supplementary Figure 4). (**E**) Similar distributions for gDNA and cDNA traces with 100ng/ml dox added at time 0 (See I0 and t0 distribution in Supplementary Figure 4). The labeled text denoted the median and the standard error of the mean. At both induction levels, the autoregulatory (gDNA) system exhibits a tighter distribution of final equilibrium SF2 levels and response rates. The negative splicing feedback loop thus reduces cell-cell heterogeneity both in final level and dynamics. (**F**) Flow cytometry data confirms that the autoregulatory (gDNA) system exhibits lower cell-cell heterogeneity across a wide range of induction levels. As in Figure 2C, we induced gDNA and cDNA cells for >24hrs, fit the Citrine intensity with a Gaussian curve (Supplementary Figure 3) and used the standard deviation parameter from the Gaussian fit to represent the variance of ectopic SF2 level between cells. Error bars represent standard error of the mean from 9 experimental replicates.

These traces (Figure 4B, Supplementary Figure 4A) also showed that negative autoregulatory splicing reduces cell-cell heterogeneity: SF2(gDNA) cells had a tighter distribution of movie curves than SF2(cDNA) cells. To quantitatively examine cell-cell heterogeneity at different induction levels, we compared SF2(gDNA) and SF2(cDNA) curves with the distribution of four parameters extracted from each movie trace (Figure 4C): I_0_ and I_f_, characterizing the initial and final fluorescence levels respectively; t_0_, the time at which the rising signal occurred; and r, the slope of the rise. At low induction levels (Figure 4D), the standard deviation of r and I_f_ relative to their median values increased from 0.04 and 0.036 in SF2(gDNA) cells, to 0.10 and 0.046 in SF2(cDNA) cells, respectively. Similarly, at high induction levels (Figure 4E), variability in r and I_f_ increased from 0.12 and 0.026 in SF2(gDNA) cells, to 0.27 and 0.066 in SF2(cDNA) cells. We also analyzed a broader range of 4-epiTC concentrations by flow cytometry (Supplementary Figure 3), and confirmed that negative autoregulatory splicing reduces cell-cell heterogeneity of I_f_ across a wide range of induction levels (Figure 4F).

The delay before activation, t_0_, varied systematically with induction level, decreasing from ∼13 hrs at low induction levels to ∼3 hrs at the highest induction levels (Supplementary Figure 4B, 4C). 3 hours is comparable to the total expected time required to synthesize mature SF2 protein (Milo et al., 2010). The longer 13 hour delay may reflect the bursty nature of transcription, which can produce extended intervals between transcriptional bursts (Larsson et al., 2019; Raj et al., 2006), and variable 4-epiTC/dox induction strength and absorption efficiency between cells. This could also explain why we did not observe an accelerated response time for the SF2(gDNA) circuit at low induction levels (compare Supplementary Figure 4A and Figure 4B). Notably, t_0_ did not differ between the SF2(gDNA) and SF2(cDNA) cell lines (Supplementary Figure 4B, 4C), suggesting that it is controlled by factors independent of the splicing regulatory circuit. Similarly, the heterogeneity and level of background signal, I_0_, remained the same for SF2(gDNA) and SF2(cDNA) cells regardless of induction level (Supplementary Figure 4B, 4C), suggesting that the sum of autofluorescence and promoter leakage was similar between the two circuits.

Taken together, these results demonstrate that negative autoregulatory splicing can speed response rates and reduce cell-cell heterogeneity in response rate and expression level. These effects are similar to those of other well-known negative autoregulatory feedback loops (Becskei and Serrano, 2000; Rosenfeld et al., 2007, 2002). However, negative autoregulatory splicing has a unique feature distinct from other types of negative feedback: its ability to simultaneously affect a large and variable ‘load’ of target pre-mRNAs (Figure 1).

### Negative autoregulatory splicing modulates response of SF2 level to splicing ‘load’

In principle, *SRSF1* negative autoregulatory splicing could operate in two opposite regimes that differ in their response to increased substrate (pre-mRNA) ‘load’ (Figure 5A). In ‘robust’ mode, the feedback strength would be independent of ‘load’ level. Despite the competition between distinct pre-mRNA substrates, SF2 is abundant in the cell, and SF2 production maintains a constant concentration in the cell. Alternatively, in an ‘adaptive’ mode, the feedback loop would modulate its negative autoregulatory strength, tuning SF2 expression in response to total substrate load. In this case, increased substrate load, like a ‘sponge’, would dilute the available SF2 per substrate molecule, reducing the feedback strength, generating more functional *SRSF1* isoform 1, and thereby producing more SF2 protein.

**Figure 5:**
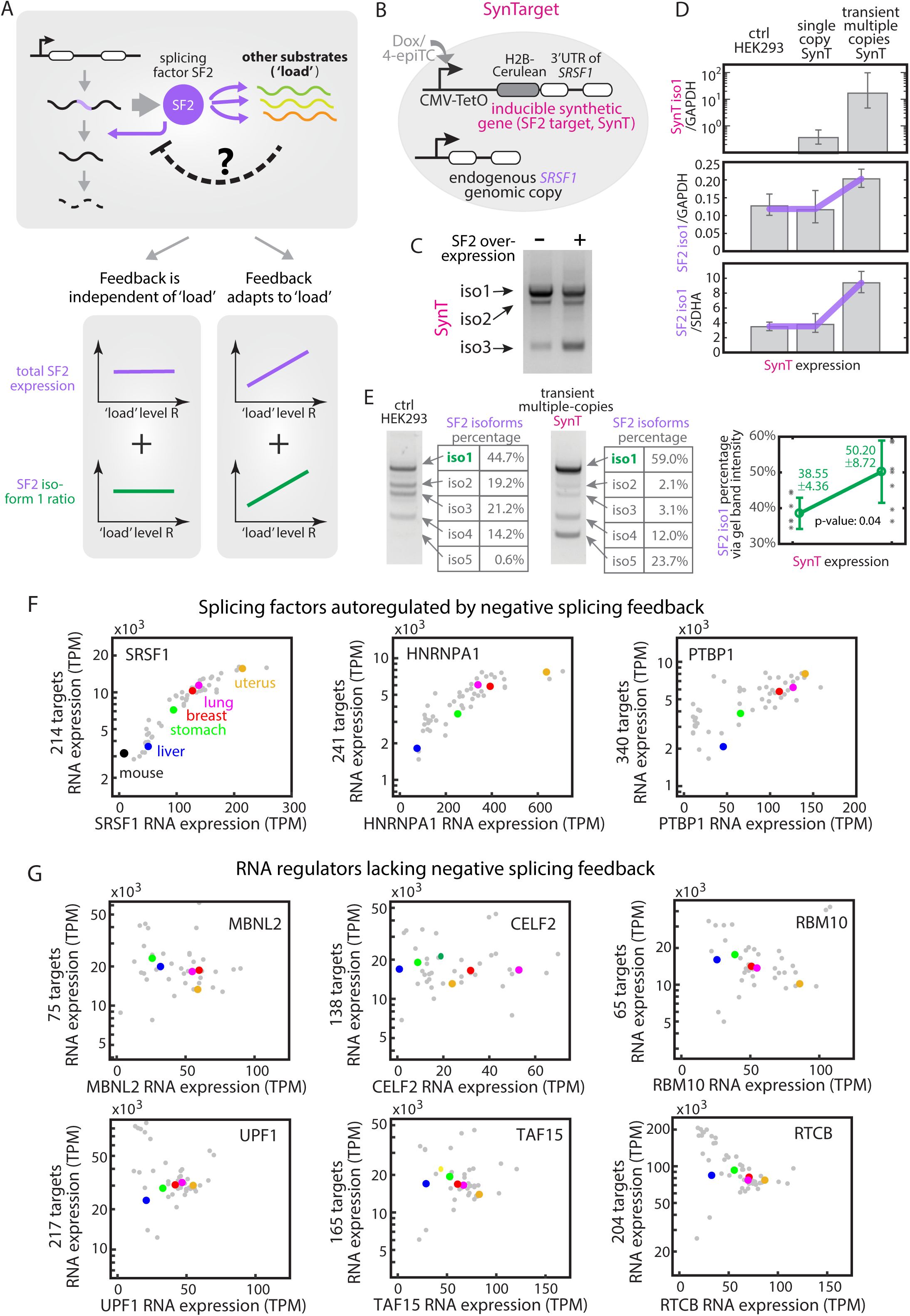
Splicing negative autoregulation modulates its feedback strength in response to variable total substrate load. (**A**) Two possible outcomes in response to total substrate ‘load’. (Left) ‘robust feedback’ scheme: the total SF2 level (purple line) and its splicing pattern (green line) remain constant across a broad range of substrate levels. The amount of SF2 involved in negative feedback is independent of ‘load’ level. (Right) ‘adaptive feedback’ scheme: more SF2 is produced (purple and green curves) via weakening negative autoregulatory splicing (i.e. dashed negative arrow in the top grey box), as increased substrate ‘load’ titrates away available SF2 in the cell. (**B**) The inducible synthetic SF2 target (SynTarget, short as SynT) cell line contains H2B-Cerulean fused with the spliceable 3’UTR of SRSF1. This synthetic gene is expressed under a Tet-On CMV promoter and stably integrated at the Flp-In locus in a T-REx HEK293 cell line. (**C**) SynT is a splicing target of SF2. We used RT-PCR and gel-imaged 3 isoforms of SynT cells with 100 ng/ml dox (left lane) and of SynT cells with transiently transfected SF2(cDNA) plasmid in 100 ng/ml dox (right lane). We found that SF2 overexpression promotes the splicing of SynT, increasing the expression of short isoforms. (**D**) We induced SynT at different levels and quantified the concentration of SynT isoform 1 (top row) and the functional SF2 isoform 1 (bottom two rows) by RT-qPCR (see qPCR qualification in Supplementary Figure 6). We found SF2 levels remained unchanged by expression from a single copy of SynT (middle column, by inducing the stably integrated SynTarget with 100ng/ml dox), but increased ∼50% when multiple SynT copies were induced in the same cell (right column, by transiently transfecting SynT plasmid with 100ng/ml dox). qPCR results were verified by normalizing to two house-keeping genes, GAPDH and SDHA, respectively. Purple solid lines are guides to the eye matching the scheme in (A). The data represents the exponential of logarithmic mean of normalized qPCR reads. Error bars represent the minimum and maximum values over 3-10 experimental replicates. (**E**) SF2 splice isoform pattern changes in response to increased total substrates. We quantified SF2 isoforms using RT-PCR and analyzed the gel band intensity by Bio-Rad ChemiDoc Image Lab 6.0 band analyzer. Two gel band examples are presented: one from HEK293 control (left), the other from transient multiple copies of SynTarget (right). Multiple copies of SynTarget trigger ∼ 30% more SF2 isoform 1 (i.e. functional unspliced isoform) through splicing. The data represents the median of gel band intensity percentage reads and error bars represent the standard deviation over 6-7 experimental replicates. Green solid lines are guides to the eye matching the scheme in (A). (**F**) The expression level of splicing factors positively correlates with their target expression across 53 human tissue types (gray dots), where 5 example tissues (uterus, lung, breast, stomach, liver) are labeled in distinct colors. The splicing factor RNA expression levels (TPM) were extracted from the GTEx database. The respective target genes are selected based on POSTAR2, specifically, the top 2% in each CLIP database. The three splicing factors, SRSF1, hnRNPA1, and PTBP1 are all autoregulated via negative splicing feedback. (**G**) The RNA regulation proteins without negative splicing feedback do not show correlative patterns as in (F).

To discriminate between these two regimes, and investigate whether and how negative autoregulatory splicing responds to perturbations of ‘load,’ we site-specifically integrated a single copy of synthetic target (SynT) of SF2 under a doxycycline(dox)-inducible CMV promoter in HEK293 cells (Figure 5B). SynT contains the spliceable 3’UTR of *SRSF1*, fused with a fluorescent protein, H2B-cerulean, at the 5’ end. It generates 3 isoforms (Supplementary Figure 5), with the same splicing junction as *SRSF1* isoforms (Supplementary Figure 1A). Transient overexpression of SF2 (cDNA) in this cell line altered the splicing pattern of SynT towards more spliced isoforms (Figure 5C), consistent with SynT acting as a SF2 target.

To determine the response of SF2 to SynT load, we need to quantify the total expression level, as well as the splicing pattern, of SF2 across different SynT induction levels (Figure 5A, bottom). The splicing pattern can be quantified using RT-PCR (as in Supplementary Figure 1B, and Figure 5C) by amplifying all isoforms at once with a single primer set targeting the 5’ and 3’ end of the gene (Materials and Methods). To quantify the total expression levels of SF2 and SynT, we used RT-qPCR. Since only SF2 isoform 1 is functional, we focused on the amount of this particular isoform. The similarity between isoform sequences limits the number of primer sets that can distinguish isoform 1 from other isoforms. Nevertheless, we identified functional priming sites and verified that the RT-qPCR responded linearly across the relevant range despite the non-optimal lengths of the RT-qPCR products (699bp and 310bp, Supplementary Figure 6). These results show that it is possible to quantify isoform expression levels using RT-qPCR.

Using this assay, we tested the response of SF2 levels and splicing patterns to ectopic expression of a single genomically integrated copy of SynT (Figure 5D, single copy SynT). Separately, to ensure effective competition with the large number of endogenous SF2 targets (Das and Krainer, 2014), we also transiently transfected multiple copies of the SynT gene that together produce ∼20-100 times more SynT (Figure 5D, transient multi-copy SynT). We observed no significant change in SF2 levels in response to the single copy perturbation. By contrast, SF2 levels increased ∼1.5-2 fold in response to transient, multi-copy SynT transfection (Figure 5D middle and lower panels).

Negative autoregulatory splicing contributed to this increased SF2 level. As shown in Figure 5E, when SynT level increased, a higher fraction of SF2 pre-mRNA remained unspliced, producing more functional isoform 1. Based on the gel band intensity, the ratio of iso1 increased about 1.30±0.08 times, consistent with the qPCR measurements in Figure 5D. This result suggests that overexpression of SF2 substrates can reduce SF2 availability, impeding the unproductive self-splicing of SF2, and thereby increasing SF2 protein synthesis.

SynT is a synthetic target gene. If SF2 similarly responds to variations in the load presented by its endogenous targets, its abundance should correlate with that of its targets. We thus obtained a list of SF2 target genes from the CLIP-seq database POSTAR2 (Zhu et al., 2019), and corresponding transcript levels (TPM) of both SF2 and its CLIP-based target genes from GTEx (www.gtexportal.org) (Supplementary Figure 7). Across 53 human tissue types (Figure 5F), SF2 expression positively correlated with that of its targets. Similarly, other splicing factors with conserved autoregulatory negative splicing feedback loops (Ni et al., 2007; Wollerton et al., 2004), such as *hnRNPA1* and *PTBP1*, also correlated with their targets (Figure 5F). In contrast, RNA binding proteins without negative splicing feedback did not present any clear correlation (Figure 5G).

Taken together, these results indicate that substrate load can modulate negative autoregulatory splicing feedback. This load-adaptive feedback scheme allows ultra-conserved splicing factors to adapt their own protein expression levels to variable substrate levels across diverse tissues and species (Hanamura et al., 1998; Lareau et al., 2007; Zahler et al., 1992).

### A mathematical model explains the dynamics and function of negative autoregulatory splicing

Having shown experimentally that negative autoregulatory splicing accelerates SF2 response times and enables their adaptation to substrate load, we sought to understand how these features arise from the autoregulatory architecture. We developed a mathematical model describing the dynamics of SF2 with and without feedback, and fit the model to SF2 dynamics observed in time-lapse movies of the SF2(gDNA) and SF2(cDNA) cell lines.

We formulated differential equations describing the dynamics of unspliced pre-mRNA *u*, functional mRNA isoform *m*, and protein *p* from the endogenous (subscript en) and ectopic (subscript ec) copies (Figure 6A). For both sets of equations, we assumed the same constant production rate (α) for either of the two ectopic constructs, controlling production of either *m*_ec_ for SF2(cDNA) or *u*_ec_ for SF2(gDNA). We also assumed that translation of *p* is linearly related to the corresponding mRNA concentration *m*, at rate γ, and that all species undergo first-order degradation with rate constant β.

**Figure 6:**
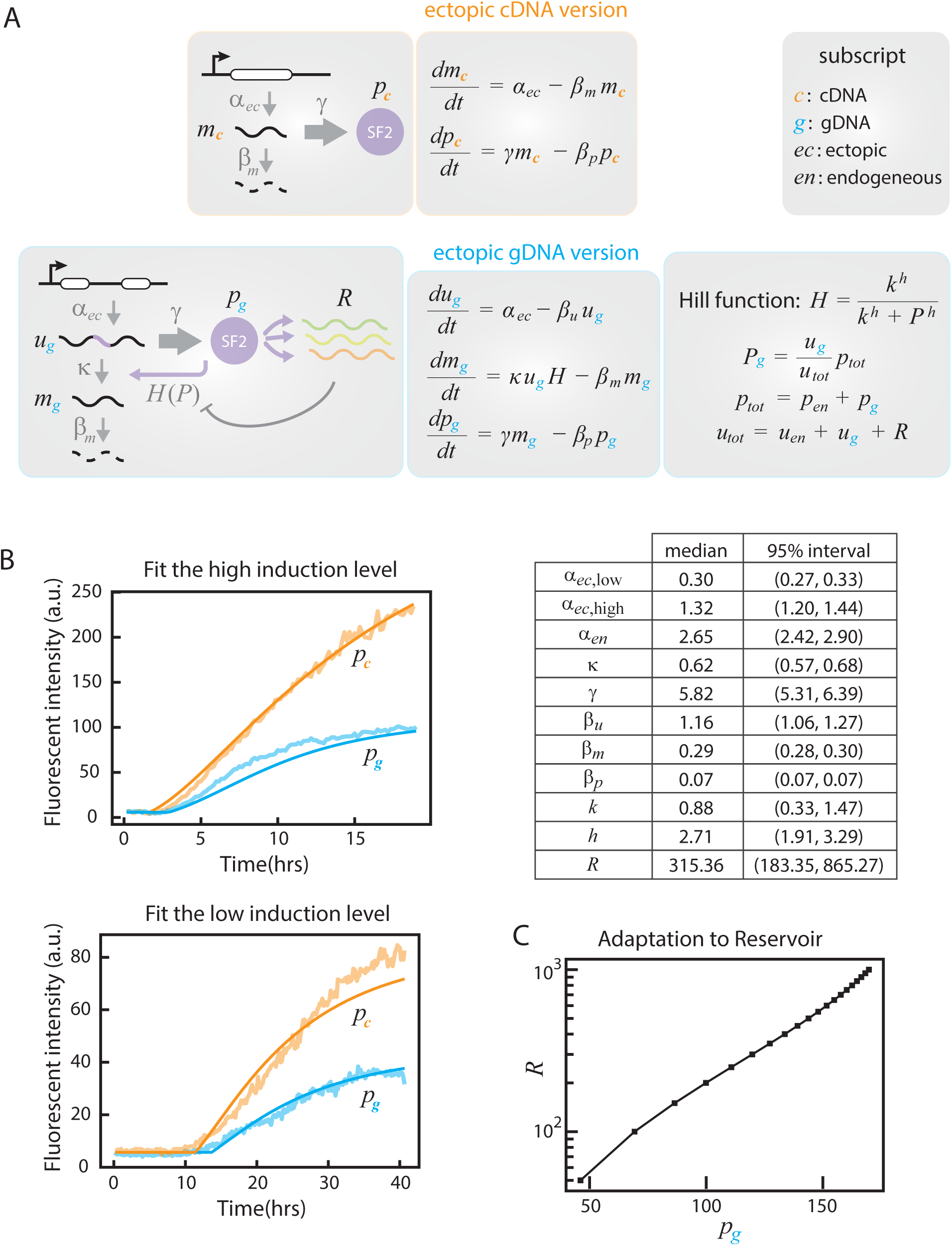
A minimal mathematical model describes splicing regulation with and without negative feedback. (**A**) Differential equation sets for the regulation of ectopic SF2(cDNA) (without feedback) and SF2(gDNA) (with feedback) cell lines. Unspliced pre-mRNA, functional mRNA and SF2 protein are labeled as *u, m* and *p*, respectively. Subscripts are defined at left. As each ectopic copy is stably integrated into the same genomic locus of HEK293 cells, they share the same transcription rate (*α*), translation rate (*γ*), degradation rate (*β*) between cell lines. Note that gDNA system incorporates a Hill function *H(P)* to represent the dependence of splicing activity on SF2 protein. For details, see Materials and Methods. (**B**) Fits of median time-lapse traces from Figure 4B and Supplementary Figure 4A to the equations in A. Fitted curves are shown with smooth solid lines. Best fit parameter estimates are shown in the table and Supplementary Figure 8. (**C**) Negative splicing autoregulation adapts to ‘load’. Using the fitted parameters, the model predicts a positive correlation between target load and SF2 level, *p*, similar to experimental observations (Figure 5).

We define two sets of equations, with and without feedback. In the case of SF2(cDNA), the functional isoform *m*_ec_ is not subject to negative autoregulatory splicing. Since the endogenous components do not have any regulatory effect, modeling the dynamics of *m*_ec_ and *p*_ec_ is sufficient (Figure 6A, ectopic cDNA version, Materials and Methods). In contrast, the SF2(gDNA) cell line features negative feedback impacting the splicing of *u*_ec_ to *m*_ec_ via both the endogenous and ectopic SF2 protein (Figure 6A, ectopic gDNA version and endogenous copy, Materials and Methods). Since SF2 also acts on many other target genes, collectively denoted *R*, we reasoned that the level of SF2 available for autoregulatory splicing would be reduced by competition with *R*. Therefore, we modeled negative feedback by a Hill function *H*(*p*), where the effective protein level acting on *SRSF1* transcripts depends on the abundance of those transcripts relative to those of the reservoir. The resulting differential equations are summarized in Figure 6A; more details on the derivation are given in Materials and Methods.

We then fit the model parameters to the averaged traces in Figure 4B using a Bayesian inference framework (Figure 6B, Materials and Methods). The data comprise ectopic SF2 protein levels for the SF2(cDNA) and SF2(gDNA) lines at both low and high induction levels (Figure 4B and Supplementary Figure 4). Shared biochemical parameters (for instance, translation and degradation rates) are fit jointly across all conditions. Parameters specific to the negative feedback case are: *κ*, the efficiency of splicing to the unproductive isoform; *k*, the substrate concentration that produces half-maximum splicing activity; *h*, the Hill coefficient; and *R*, the reservoir level (Materials and Methods). Only the ectopic production rate was allowed to change between low and high induction levels.

We computed probability distributions for each parameter (Supplementary Figure 8) and identified median values and confidence intervals (Figure 6B Table). These values were approximately consistent with independent parameter estimates. In particular, the ratio between production rates of the ectopic and endogenous *SRSF1* copy in SF2(gDNA) cells (i.e. α_*ec*_ versus α_*en*_) matched that obtained from the western blot in Supplementary Figure 2. Similarly, the fit degradation rate values generate half-lives of about 35 minutes for *u*, 2.4 hours for *m*, and 10 hours for *p*, broadly consistent with values in other studies (Milo and Phillips, 2015; Moulton et al., 2014). Finally, the fitted value of *κ* suggests that 62% of transcripts are spliced to isoform 1, generally consistent with the percentage seen in multi-copy SynT transfection (Figure 5E).

The fitted parameters can provide deeper insight into the biology of SF2 regulation. The Hill coefficient values, *h*∼2-3, suggests ultra-sensitivity of SF2 activity. The estimated reservoir level *R* of about 300 suggests approximately several hundred additional SF2 target transcripts per SF2 transcript in this cell type, also consistent with typical values in a variety of cell types (Figure 5F). Finally, using the model to compute the steady-state ectopic protein level as a function of *R* reveals that SF2 level adapts to target load (Figure 6C), consistent with experimental results (Figure 5).

## Discussion

Negative autoregulation is a prominent feature of many splicing regulatory systems. To understand its functional impact, we constructed three HEK293 cell lines, with inducible gDNA (autoregulated), cDNA (unregulated), and SynT (synthetic SF2 targets) and quantitatively investigated the dynamic function of negative autoregulatory SF2 splicing in individual cells. By combining deep-learning based single cell movie analysis (Figure 3) with flow cytometry, RT-qPCR and RT-PCR, we found negative autoregulatory splicing can stabilize SF2 levels independent from its own transcription strength, reduce cell-cell heterogeneity in both SF2 levels and response times (Figure 4), and maintain constant SF2 activity despite changes in target load (Figure 5). Although we focused on SF2, the approach presented here can be extended to study other splicing factors.

A particularly interesting aspect of negative splicing autoregulation compared to other levels of negative autoregulation is its ability to homeostatically control splicing activity in response to changes in total substrate load. Thus, even though both SF2 and SynT overexpression increases absolute SF2 levels, they produce distinct, and opposite, effects on *SRSF1* splicing patterns (Supplementary Figure 1B, Figure 5D), with more spliced SF2 isoforms produced when SF2(cDNA) was overexpressed (Supplementary Figure 1B), and more unspliced SF2 isoforms when SynT was overexpressed (Figure 5E). These results confirm that the cell maintains sub-saturating SF2 levels. To maintain a constant effective concentration, negative autoregulatory splicing buffers SF2 activity against variations in both absolute SF2 levels (Figure 2, 4) as well as target levels (Figure 5).

In addition to the experimental data, mathematical modeling provided additional insights into SF2 autoregulation. First, it shows that experimentally observed effects can arise from basic aspects of splicing and transcriptional regulation more generally, even in the absence of additional, complex mechanisms. However, the model does not rule out more complex mechanisms. Further molecular analysis will be necessary to fully characterize mechanistic aspects of autoregulation. Second, the model provides a unified understanding of the sometimes counterintuitive responses of SF2 to different perturbations driven by feedback and self-splicing. Third, the model identified biochemical parameters that recapitulate experimental observations and can help to inform a more quantitative understanding of splicing dynamics. Finally, the fitted parameter values suggest that SF2 activity may exhibit ultrasensitive dependence of activity on its own concentration.

This analysis does not explicitly incorporate the non-uniform spatial distribution of SF2 in the nucleus. SF2 concentrates in interchromatin granule clusters called speckles (Lamond and Spector, 2003; Misteli et al., 1997) that exhibit dynamic structures (Misteli et al., 1997). Recent work suggests that the spatial distribution of splicing factors correlates with active transcription hubs in the nucleus (Ding and Elowitz, 2019; Quinodoz et al., 2018). Because most splicing occurs co-transcriptionally (Bentley, 2014; Das et al., 2007; Rosonina and Blencowe, 2002), negative autoregulatory splicing should, in principle, feedback on the local SF2 concentration within a sub-nuclear neighborhood, rather than the global concentration averaged over the nucleus as a whole. It thus remains unclear how splicing factors balance their local and global concentrations in the nucleus. In the future, it will be interesting to develop a more complete analysis of negative autoregulatory splicing that includes SF2 subcellular spatial distribution and may provide an integrated view of how cells maintain constant effective SF2 concentrations despite heterogeneity in their subnuclear spatial distributions.

## Materials and Methods

### Cell line construction

Flp-In™ T-Rex™ HEK293 cells (Life Technologies, we did not test for mycoplasm) were cultured following the manufacturer’s protocol. For transfection, the cells were pre-plated in 24-well plates with 80% confluency. We added 800–1000 ng of plasmid (SF2(gDNA) or SF2(cDNA) or SynT) using the Lipofectamine LTX plasmid transfection reagent (ThermoFisher Scientific) and changed the culture media to Opti-MEM™ Reduced Serum Medium (ThermoFisher Scientific). Cells were left in the incubator overnight, then trypsinized (using 0.25% Trypsin-EDTA (Thermo Fisher Scientific)) into new 6-well plates with complete culture media the next day. These cells were then cultured for 1–2 weeks with 100 ug/ml Hygromycin. The surviving transfected cells were subcloned by limiting dilution.

### Time-lapse microscopy imaging and data analysis

24-well 10 mm diameter glass No. 1.5 coverslip plates (MatTek Corp.) were coated with 5 ug/ml Human Fibronectin (Oxford Biomedical Research, Rochester Hills, MI) in PBS buffer for 1hr at room temperature. Fibronectin was then aspirated, and 4,000 - 10,000 cells were plated in the coated 24-well plate with complete cell media. The plate was manually swayed (Hui and Bhatia, 2007) to uniformly spread the cells, and left in the incubator for 2-3 hrs before imaging. Details of microscopy have been described previously (Nandagopal et al., 2019). For each movie, 40-60 stage positions were picked manually, and YFP and differential interference contrast (DIC) images were acquired every 10 mins with an Olympus 20x objective using automated acquisition software (Metamorph, Molecular Devices, San Jose, CA). The details of cell tracking algorithm was in (Bintu et al., 2016; Singer et al., 2014).

### Flow cytometry

Experimental procedures and data analysis for flow cytometry have been described previously (Nandagopal et al., 2019).

### Transient transfection

Cells were plated at 50% confluency in 24-well plates and grown to 80% confluency overnight. We added 800–1000 ng of plasmid (SF2(gDNA) or SF2(cDNA) or SynT) using the Lipofectamine LTX plasmid transfection reagent (ThermoFisher Scientific) and changed the culture media to Opti-MEM™ Reduced Serum Medium (ThermoFisher Scientific). Cells were left in the incubator for 6/10/20 hrs, then trypsinized (by 0.25% Trypsin-EDTA (Thermo Fisher Scientific)) off the plates and resuspended in PBS (Phosphate-Buffered Saline buffer, Thermo Fisher Scientific). These cells were centrifuged, and the pellet washed 3x with PBS buffer to remove Trypsin. The final cell pellets were either dissolved for RNA extraction, or frozen and stored at -80C.

### RT-PCR

We extracted total RNA using the RNeasy Mini Kit (Qiagen), and used 500 ng - 1 ug RNA to make cDNA using the iScript cDNA Synthesis Kit (Bio-Rad Laboratories, Hercules, CA). 0.1 of cDNA (i.e. 1ul after diluting cDNA 10x) was used in the PCR reaction, using AccuPrime™ Pfx SuperMix (Thermo Fisher Scientific) with an annealing temperature at 62.5 degree for 35 cycles. PCR primers are listed in Supplementary Table 1. 3-8 ul of the PCR product was then run on a 1% Agarose gel. Gel band intensity was analyzed by Bio-Rad ChemiDoc Image Lab 6.0 band analyzer.

### PT-qPCR

RNA was extracted as in RT-PCR. We then used 1 ug RNA to make cDNA using the SuperScript™ III First-Strand Synthesis kit (Thermo Fisher Scientific) with all gene-specific primers (*SRSF1*, SynT, *GAPDH*, and *SDHA* isoform1 gene-specific primers). We used gene-specific cDNA for qPCR, rather than random cDNA, to minimize the influence between isoforms with similar sequences. Experimental procedures and data analysis of qPCR were performed as described previously (Nandagopal et al., 2019). All primers are listed in Supplementary Table 1.

### Western blot

Frozen or fresh cell pellets (with 10^6^ cells) were denatured using 200ul SDS loading buffer (1x sodium dodecyl sulfate (Sigma-Aldrich), 1x protease inhibitor, 4mM EDTA) and heated for 5mins at 68 °C in a water bath. The heated cells were then centrifuged at 55,000 rpm for 1hr at 4 °C. 30 ul of the supernatant was loaded onto a NuPAGE™ 4-12% Bis-Tris Protein Gel (Thermo Fisher Scientific) and transferred using iBlot™ Transfer Stack to nitrocellulose (Thermo Fisher Scientific) following the manufacturer’s protocol. The blot was then blocked in 1xTBST, 5% dry milk, 2% BSA for 1 hr at room temperature, followed by overnight incubation at 4 °C with the primary antibody, anti-SF2 antibody (ab133689) at 1:1000 and anti-GFP (ab1218) at 1:2000, together. The next day, the blot was washed with 1x TBST three times, then incubated with anti-rabbit and anti-mouse HRP conjugated secondary antibody (GE Healthcare Life Sciences) at 1:2000 for 2hrs at 4°C. The blot was then washed with 1x TBST at room temperature for 1hr and five more times for 3 mins. Gel bands were detected using SuperSignal West Femto Chemiluminescent Substrate (Thermo Fisher Scientific) and analyzed using a Bio-Rad ChemiDoc Image Lab 6.0 band analyzer.

### Mathematical model with and without negative splicing feedback

For the cDNA construct, ectopic mRNA transcripts *m*_ec_ are generated with some production rate *α* _ec_, while protein *p*_ec_ is translated from mRNA with rate *γ*. Both species are degraded by first-order kinetics with rate constants *β*_*m*_ and *β*_*p*_ respectively. Therefore, their dynamics are described by the following set of differential equations:

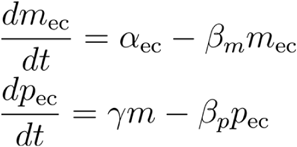

For the gDNA construct with negative autoregulation of splicing, we must consider not only mRNA and protein levels *m* and *p* but also unspliced pre-mRNA levels *u*. Here, ectopic *u*_ec_ is generated with production rate *α* _ec_ and endogenous transcripts *u*_en_ with rate *α* _en_. These transcripts are spliced productively to *m*_ec_ and *m*_en_ with rate *κ*. This splicing is regulated by negative feedback from the effective SF2 protein level *P*, which we describe phenomenologically by a Hill function

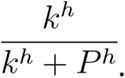

The total protein level comprises both endogenous and ectopic SF2, or *p*_tot_ = *p*_en_ + *p*_ec_. However, SF2 acts not only on ectopic and endogenous *SRSF1* transcripts but also on other target RNAs, or a “reservoir” *R*. Therefore, the effective protein level for any given target is not the total amount. SF2 not only regulates the ectopic and endogenous *SRSF1* transcripts but also acts on other target RNAs, or a “reservoir” *R*. We assume that the effective protein level for a given target is determined by the relative proportion of the target RNA. Defining the total level of unspliced targets by *u*_tot_ = *u*_en_ + *u*_ec_ + *R*, the effective protein level acting on endogenous *SRSF1* is

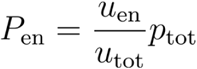

and similar for ectopic transcripts. Pre-mRNAs undergo first-order decay with rate constant *β* _*u*_; other biochemical parameters - translation rate as well as mRNA and protein degradation rates - are shared with the no-feedback model. The dynamics are given by the following set of differential equations:

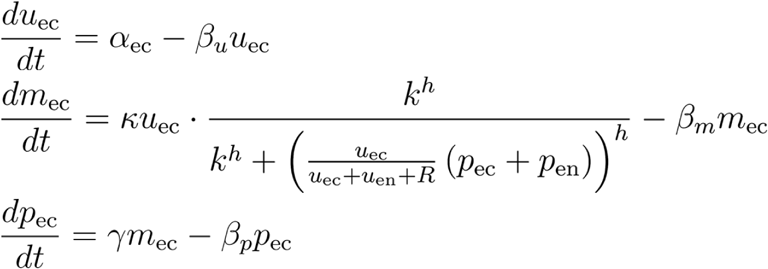

The equations for the endogenous species are analogously defined:

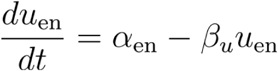

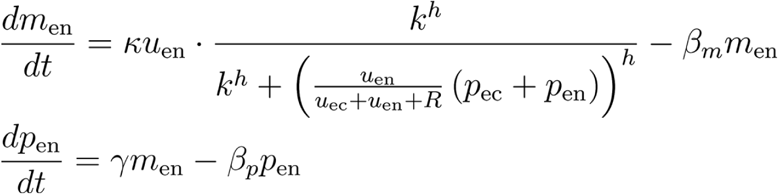

### Model parameters

We fit this model to the averaged movie curves of Figure 4, representing SF2 protein levels with and without feedback for both low and high induction levels. For the no-feedback case, the relevant model parameters include the transcription rates of the ectopic construct with low 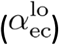 or high 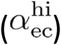 induction, the translation rate (*γ*), and the degradation rates of mRNA (*β* _*m*_) and protein (*β* _*p*_). Since we measure only the ectopic SF2 protein, we do not need to model the endogenous SF2 here.

Once we introduce regulation at the level of splicing, we must consider the ectopic as well as the endogenous SF2. We also include the parameters describing the feedback: *κ*, the ratio of full transcript spliced to the productive isoform; *k*, the activation coefficient (substrate concentration of half-maximum activity); *h*, the Hill coefficient; and *R*, the reservoir level.

In addition to these key model parameters, we fit the starting levels of endogenous SF2 species *u*_0_, *m*_0_, and *p*_0_. Based on the analysis of Figure 4, we account for time delays between induction and cellular response, considering two values *t*_low_ and *t*_high_ for the low- and high-induction conditions. We also account for background noise by setting a floor *bg* on the predicted output.

### Parameter fitting

To determine the model parameters, we performed Bayesian inference using Stan (Carpenter et al., 2017) to sample the probability distributions for each parameter given the experimental data. Bayesian statistics requires specifying priors, or the probability distributions for each parameter based on prior knowledge, and the likelihood, or the probability of observing the data given a set of parameters. Together, they allow inference of the posterior, or the probability distributions for each parameter after observing the experimental data.

For the likelihood, we assumed that experimental values lie in a Gaussian distribution centered at the theoretical value with a standard deviation *σ*. For the priors, we assumed relatively broad distributions based on biological knowledge and included applicable constraints. These prior distributions for each parameter are summarized in Supplementary Table 2. All parameters are constrained to be nonnegative; any additional constraints are listed.

For these differential equations, the starting levels of each species can be fit, but the starting time is not permitted to vary in this framework. Therefore, we screened a broad range of choices of time delays, performed an initial fit for all possible combinations, and selected the values that minimized the resulting error. These initial fits were done on 4 chains with 250 iterations per chain (125 warm up, 125 sampling). For time delays, we evaluated a range of 6-9 hours in the low-induction case and a range of 0-1 hours in the high-induction case, sampling the ranges at intervals of 0.25 hours (15 minutes). To quantify error for each set of time delays, we took the median value across samples for each parameter, simulated the resulting curve, and calculated the sum of squared errors (SSE) between the theoretical and observed values. This procedure yielded an optimal set of time delays of 8.25 hours for the low-induction condition and 0.25 hours for the high-induction condition. We then reran a more extensive fitting for this set of values, using 4 chains with 2000 iterations per chain (1000 warm up, 1000 sampling). The resulting samples are the basis for all results presented in the main text.

### Data Availability Statement

All original data, python code, DNA constructs, and cell lines are available upon request.

## Acknowledgements

We thank F. Tan for providing Western blot protocol, N. Nandagopal and Y. Antebi for technical assistance, and Life Sciences Editors (Sabbi Lall) for critical feedback on the manuscript. We also thank S. Sun, A.R. Krainer, D. Sprinzak, D. Baltimore, J.G. Ojalvo, N. Wingreen for discussion and feedback on the project. F.D. was supported by a Fellowship from the Schlumberger Foundation. C.S. is supported by the NIH National Institute of General Medical Sciences training grant GM008042 and by a David Geffen Medical Scholarship. The work was funded by the Gordon and Betty Moore Foundation through Grant GBMF2809 to the Caltech Programmable Molecular Technology Initiative and the Institute for Collaborative Biotechnologies through grant W911NF-09-0001 from the U.S. Army Research Office, AWS Machine Learning Research Awards, and the Intel Corporation. The content of the information does not necessarily reflect the position or the policy of the Government, and no official endorsement should be inferred. M.B.E. is a Howard Hughes Medical Institute Investigator.

## Author Contributions

F.D. conceived of the project. F.D. and M.B.E. designed experiments. F.D. and K.C. performed experiments. F.D. analyzed data. C.S. and F.D. did mathematical modeling. F.D., C.S., and M.B.E. wrote the manuscript.

---

**Supplementary Figure 1:**
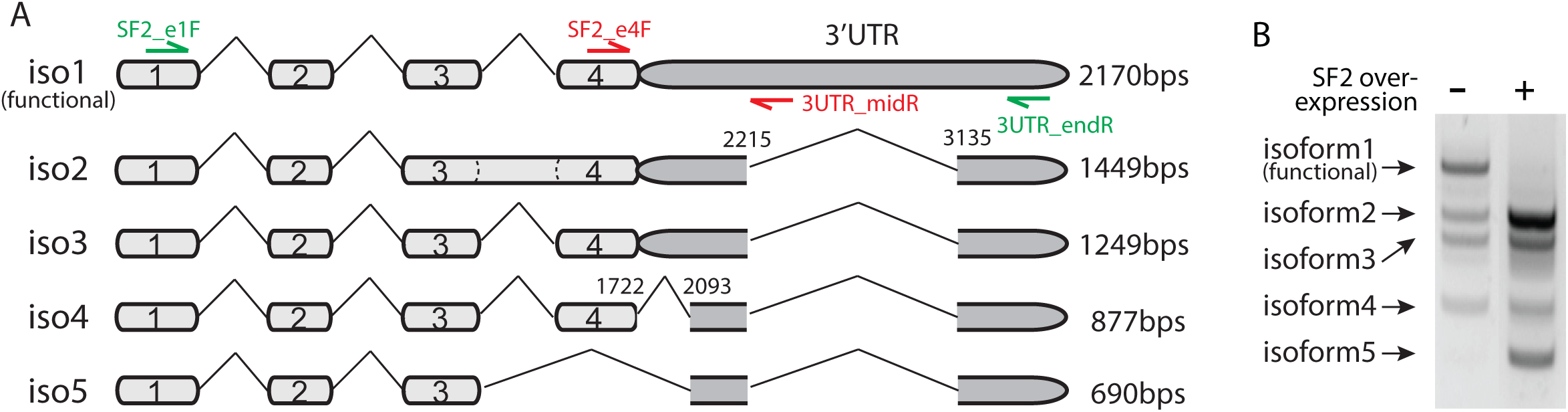
(**A**) 5 isoforms of SRSF1 and Citrine-fused SRSF1 were observed in HEK293 cells. The sequences were obtained by Laragen@ sequencing after RT-PCR (Materials and Methods) and gel extraction. Labeled primers are for RT-PCR (green) and RT-qPCR (red) respectively (Supplementary Table 1). (**B**) SF2 overexpression promotes unproductive splicing of its own gene. We used RT-PCR and gel-imaged 5 isoforms of SRSF1 of HEK293 cells (left lane) and of SF2(cDNA) cells with maximum induction level (100ng/ml dox) for 50hrs (right lane). We found that SF2 overexpression reduced the ratio of isoform 1 (i.e. the functional isoform that can be productively translated to SF2(Sun et al., 2010)), and increased expression of the other 4 unproductive isoforms.

**Supplementary Figure 2:**
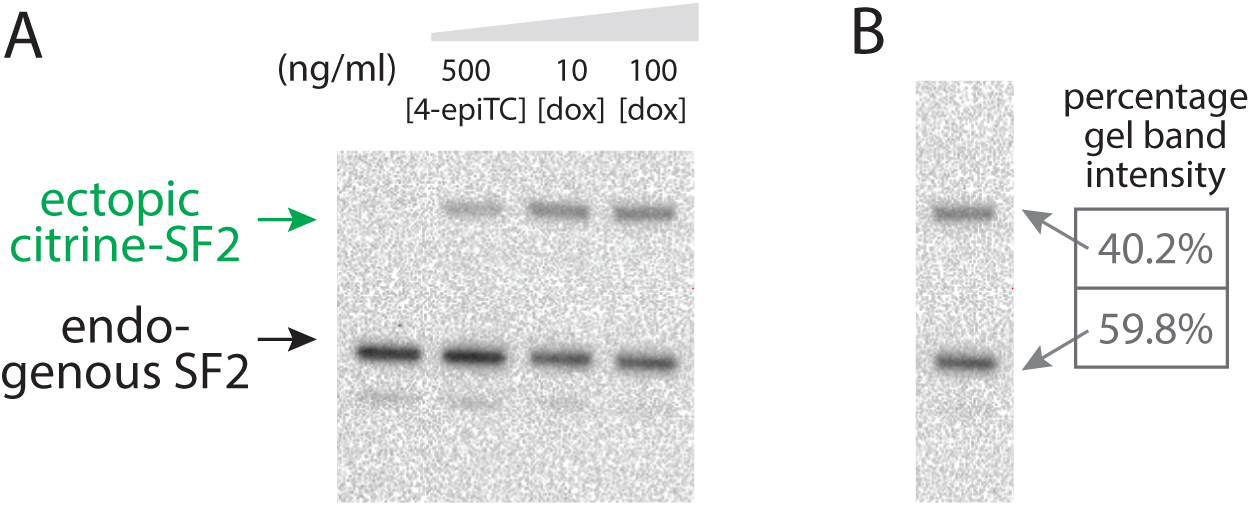
(**A**) Western blot (Materials and Methods) shows that endogenous SF2 levels decrease with increasing expression of the ectopic copy. We induced SF2(gDNA) cells at different 4-epiTC/dox concentration for 24hrs. Antibody staining details as in Figure 2B. (**B**) Endogenous SRSF1 in HEK293 cells has a high level of transcription, comparable to the ectopic CMV promoter. We analyzed the gel band intensity with highest induction level using a Bio-Rad Chemi-Doc Image Lab 6.0 band analyzer.

**Supplementary Figure 3:**
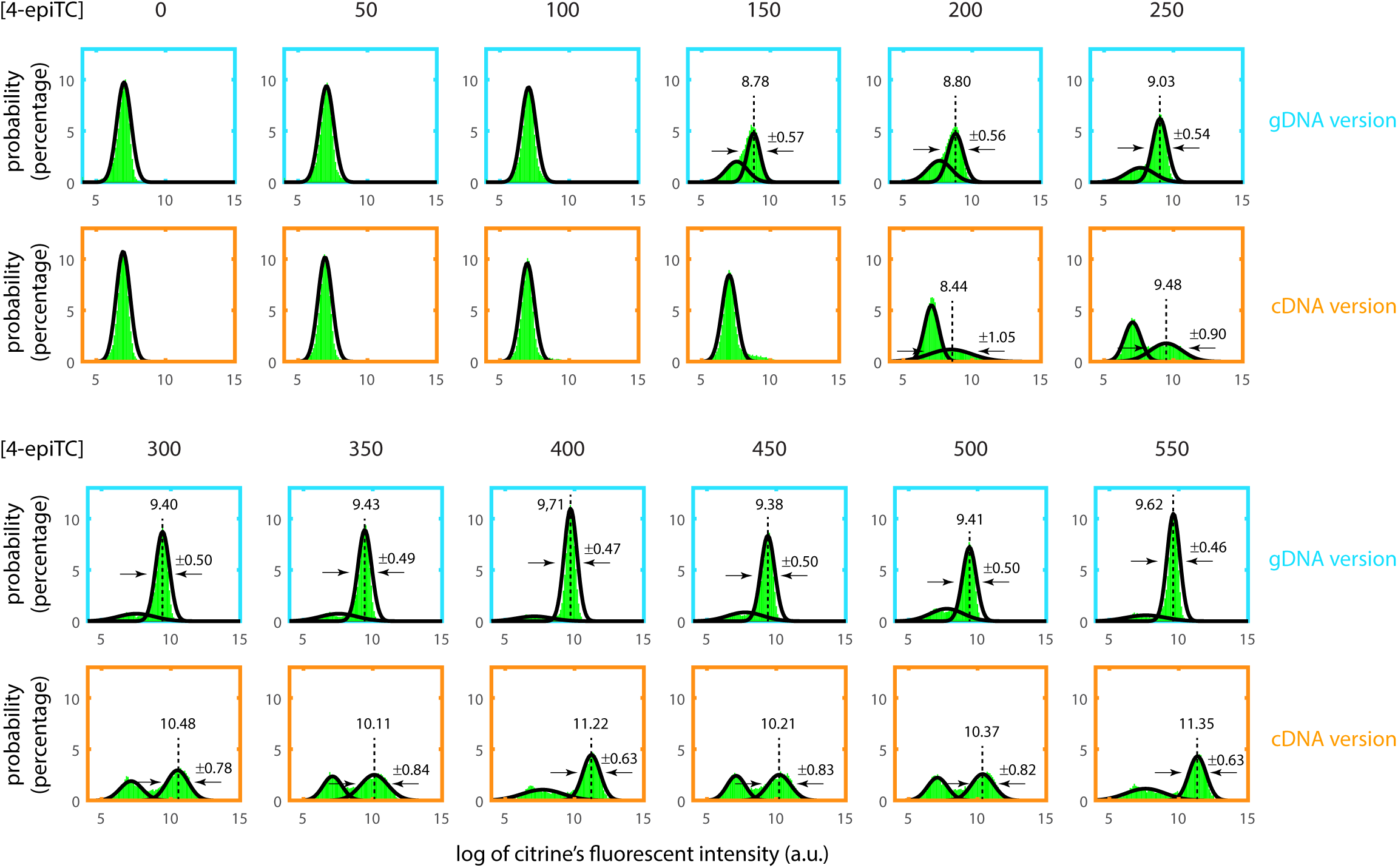
Flow cytometry data were analyzed by Gaussian fitting. One typical example of experimental replicates is shown here. The apparent bimodal distributions with short induction time (<24hrs) are probably due to cell-cell heterogeneity in 4-epiTC absorption efficiency, and stochastic transcriptional noise from the CMV promoter. This bimodality diminishes for longer induction time. To minimize the impact of this bimodal effect, we used the mean of Gaussian fitting from only fully induced cells (i.e. high peak) to represent SF2 expression in Figure 2C. The Gaussian fitting center and variance are labeled in each plot.

**Supplementary Figure 4:**
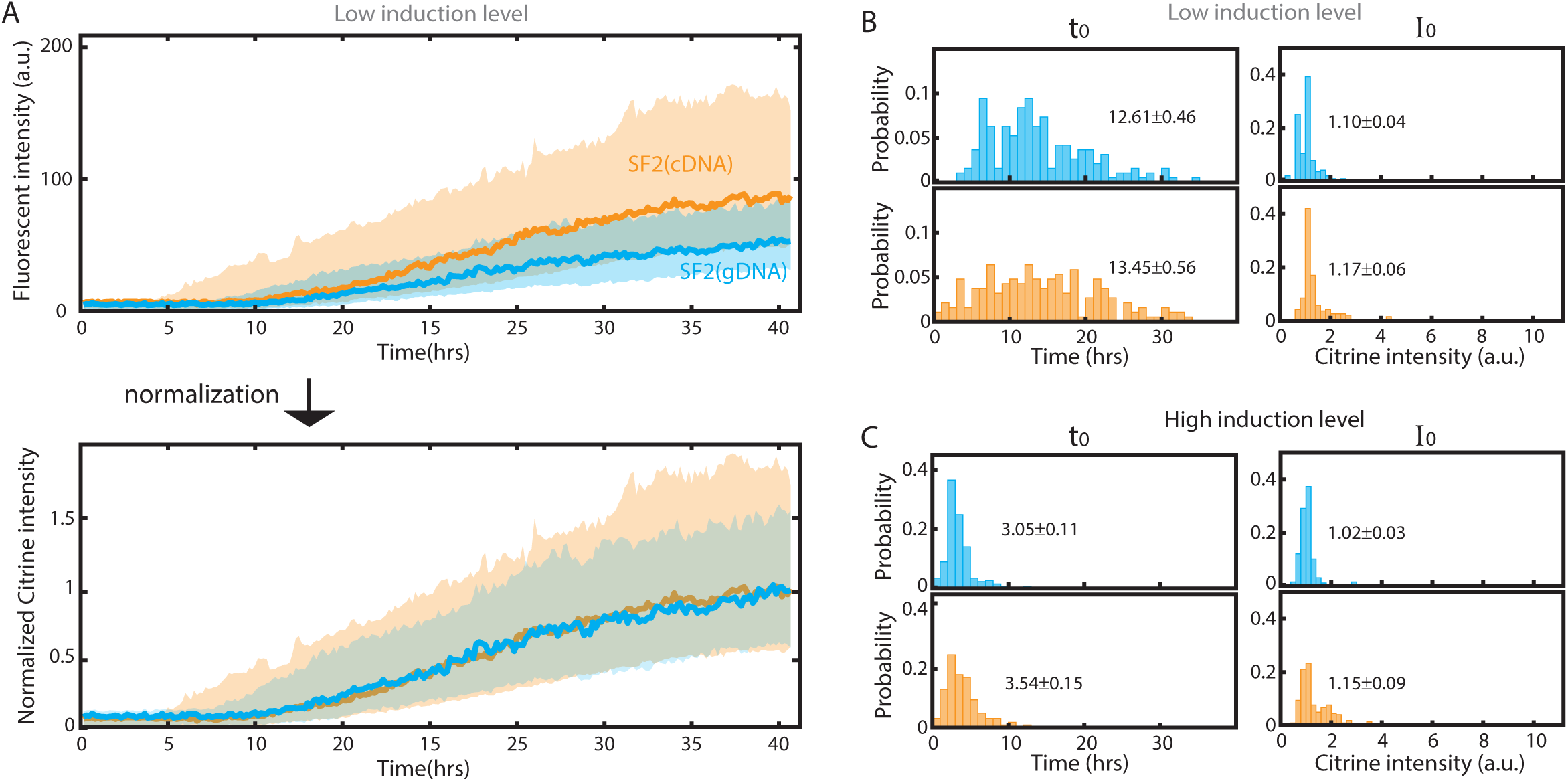
(**A**) Negative autoregulatory splicing reduces cell-cell heterogeneity in both response rate and steady-state SF2 expression at low induction level. (Top) Solid curves are the median of 191 SF2(cDNA) and 188 SF2(gDNA) single cell traces. Shading represents the standard deviation of the mean. (Bottom) The curves are normalized to final expression. (**B**) and (**C**) are distribution of fit-parameters t0 and I0 defined in Figure 4C. Color is the same as in Figure 4D and 4E.

**Supplementary Figure 5:**
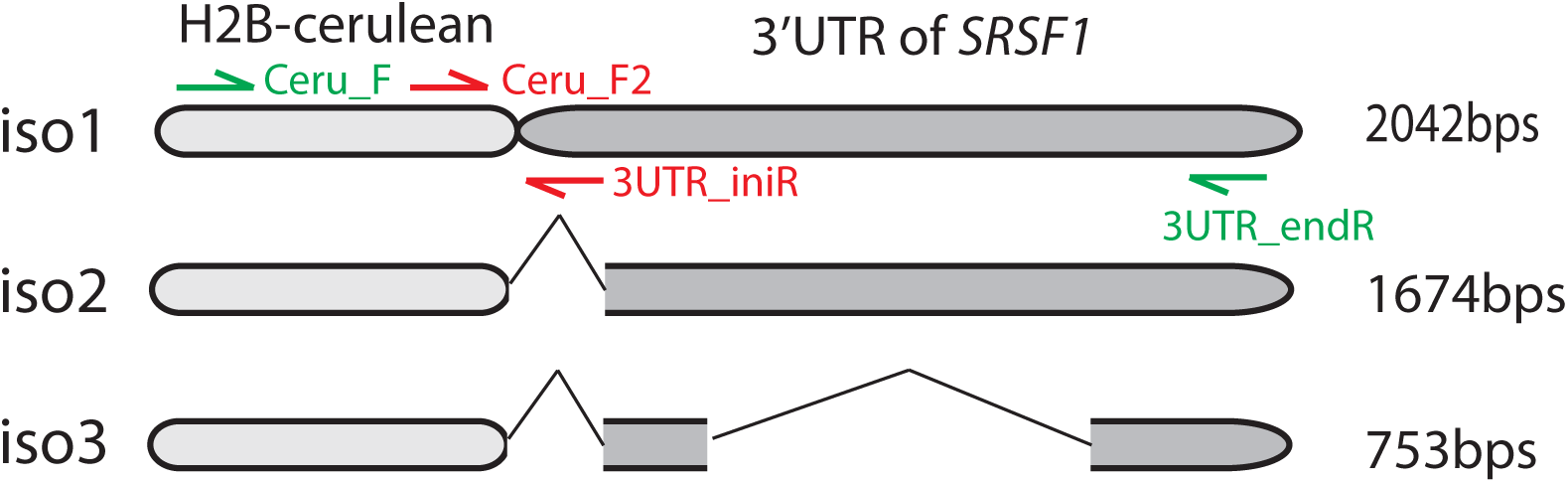
We found 3 isoforms of SynT and sequenced each isoform using Laragen@ sequencing after RT-PCR (Materials and Methods). Labeled primers are for RT-PCR (green) and RT-qPCR (red) respectively (Supplementary Table 1).

**Supplementary Figure 6:**
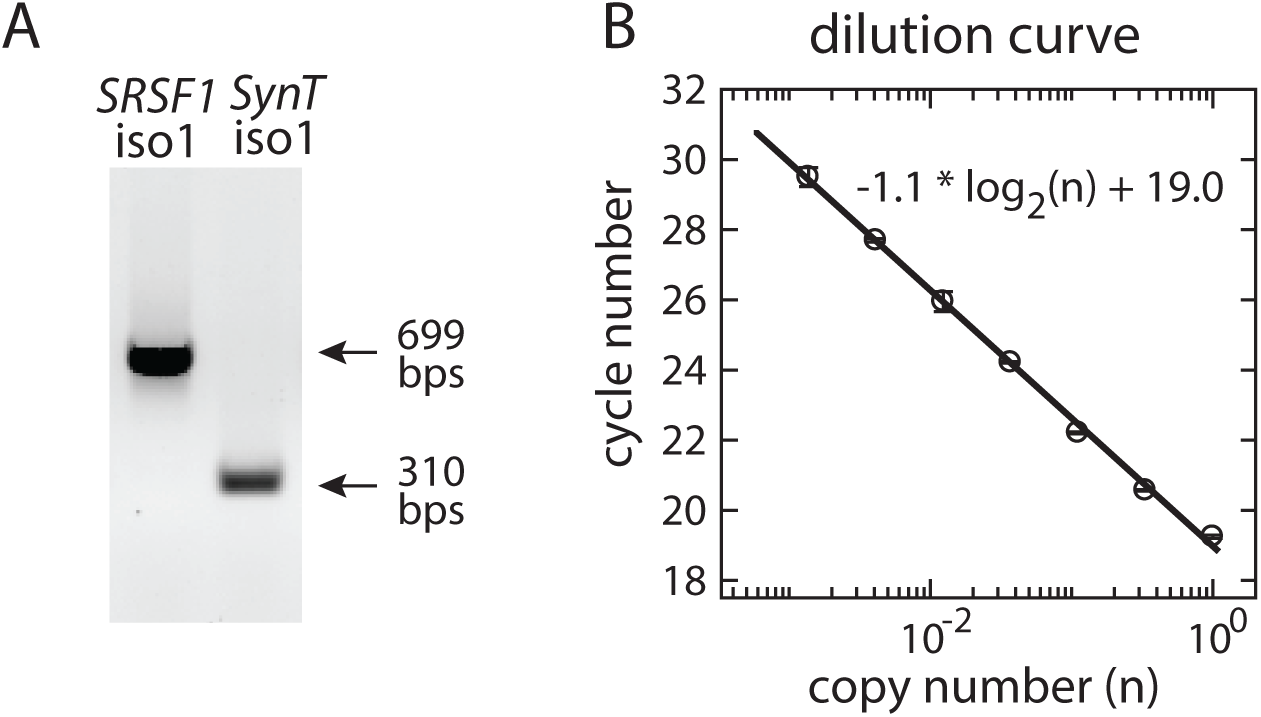
Quality control of RT-qPCR. (**A**) The products of SRSF1 isoform 1 and SynT isoform 1 RT-qPCR are 699bps and 310bps respectively. The single band indicates the specificity of qPCR amplification. (**B**) The unusual 699bps length qPCR still obeys linear rules in the dilution curve.

**Supplementary Figure 7:**
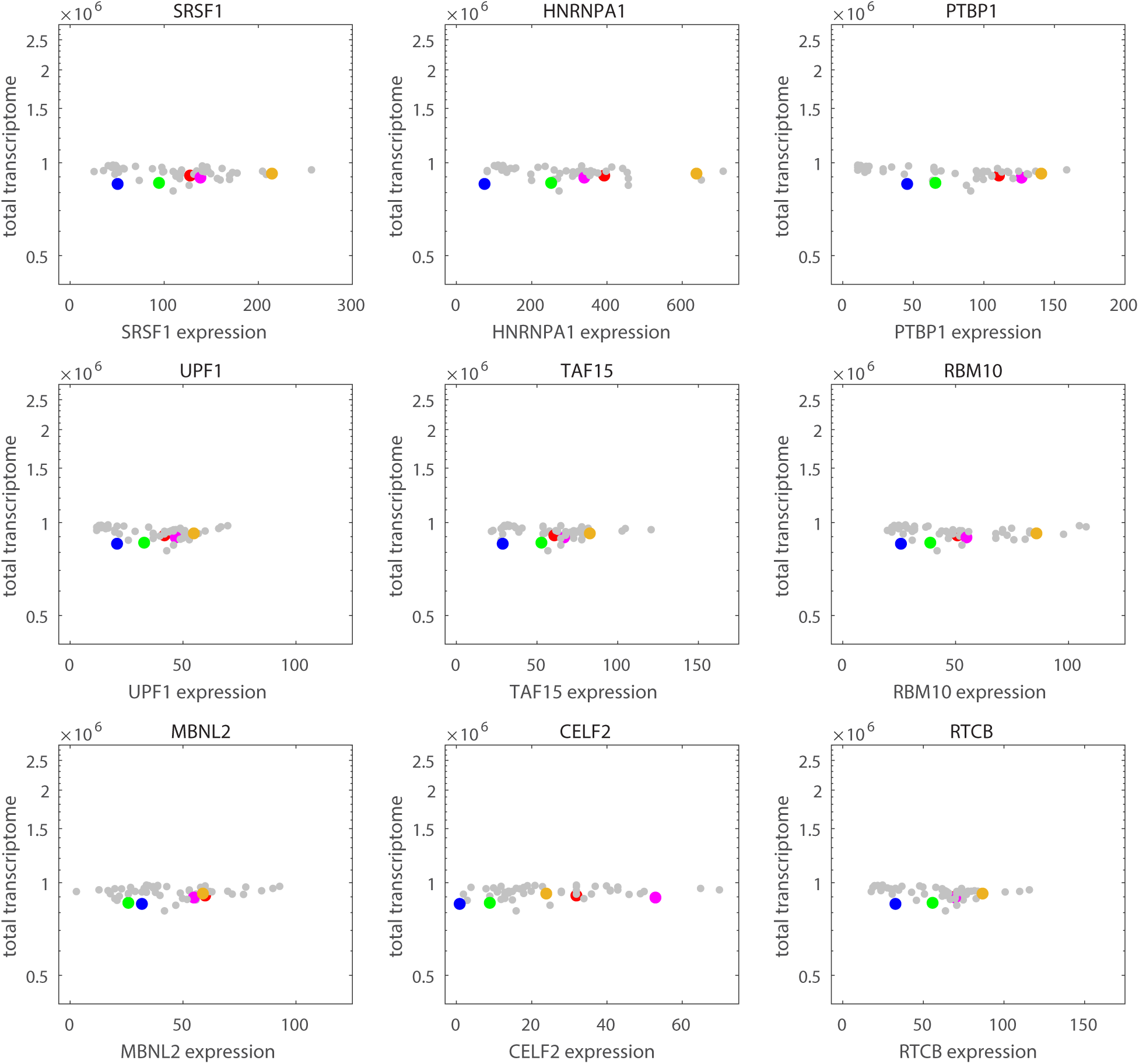
Summing up TPM of all genes (i.e. total transcriptome) from the GTEx dataset (www.gtexportal.org) shows a constant value (∼106), indicating the value is properly normalized across different tissue types.

**Supplementary Figure 8:**
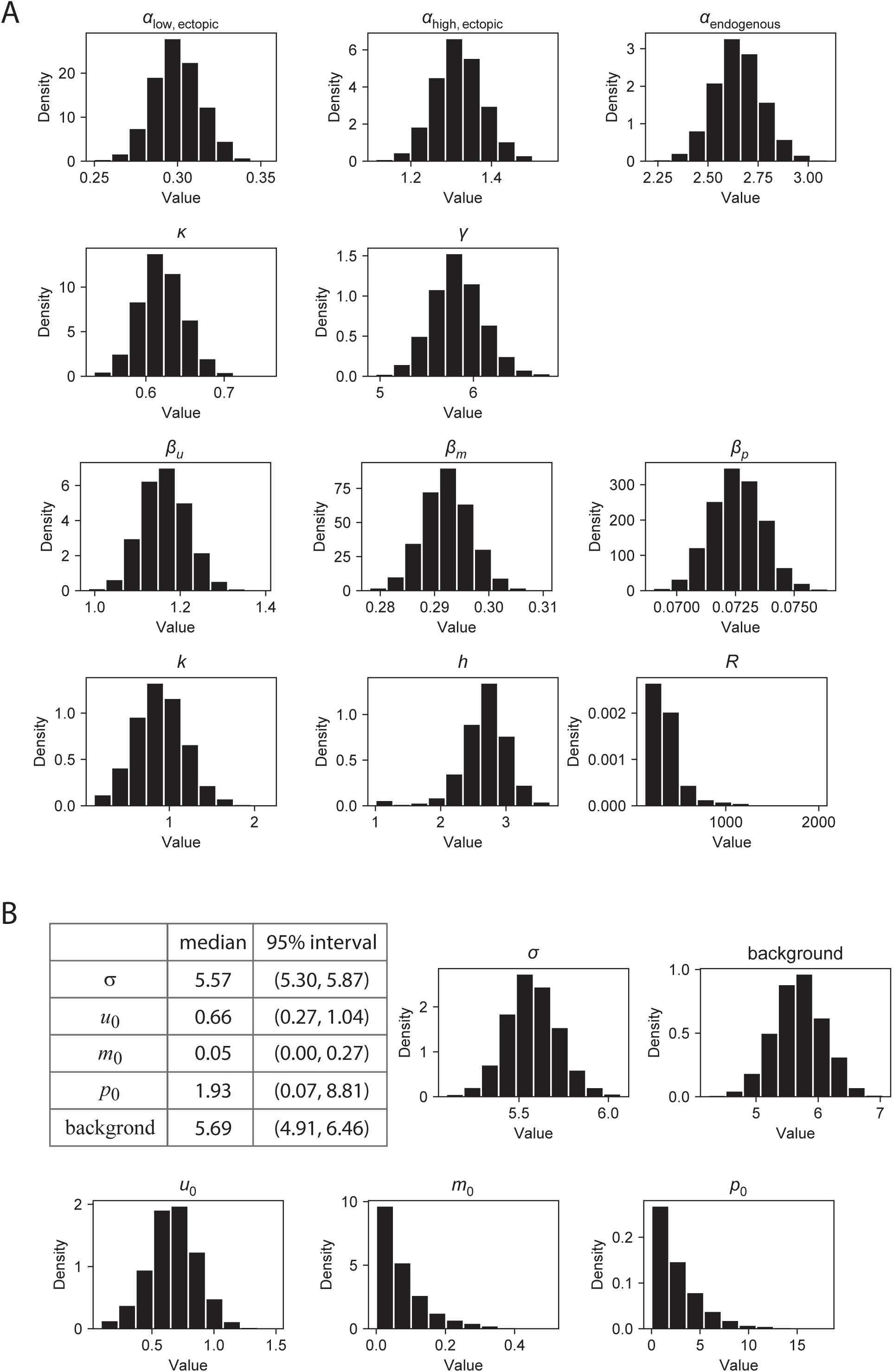
(**A**) Distribution of all parameters shown in Figure 6B Table. (**B**) The fitting estimates and distribution of all other parameters.

**Supplementary Table 1.**
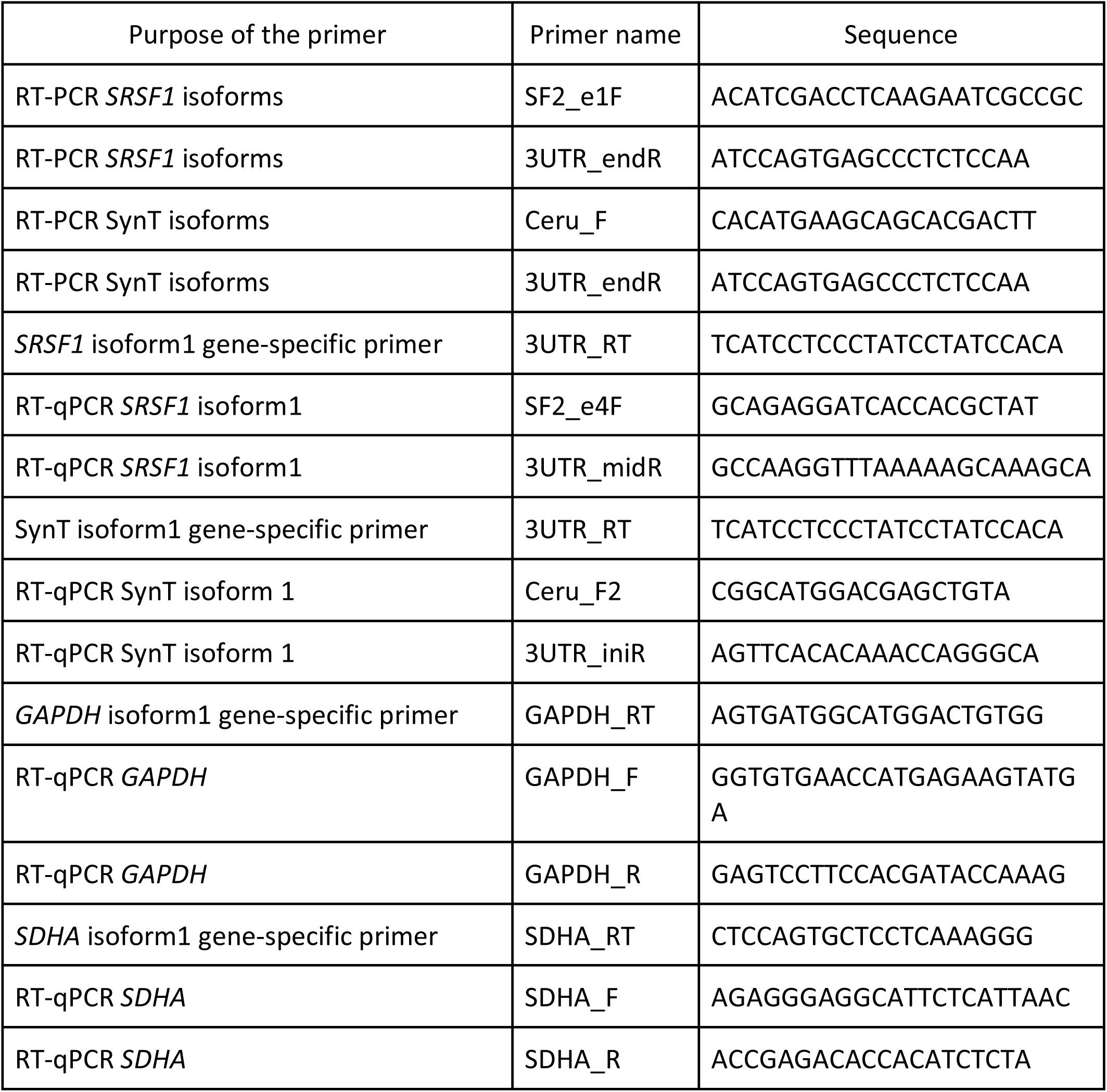

**Supplementary Table 2.**
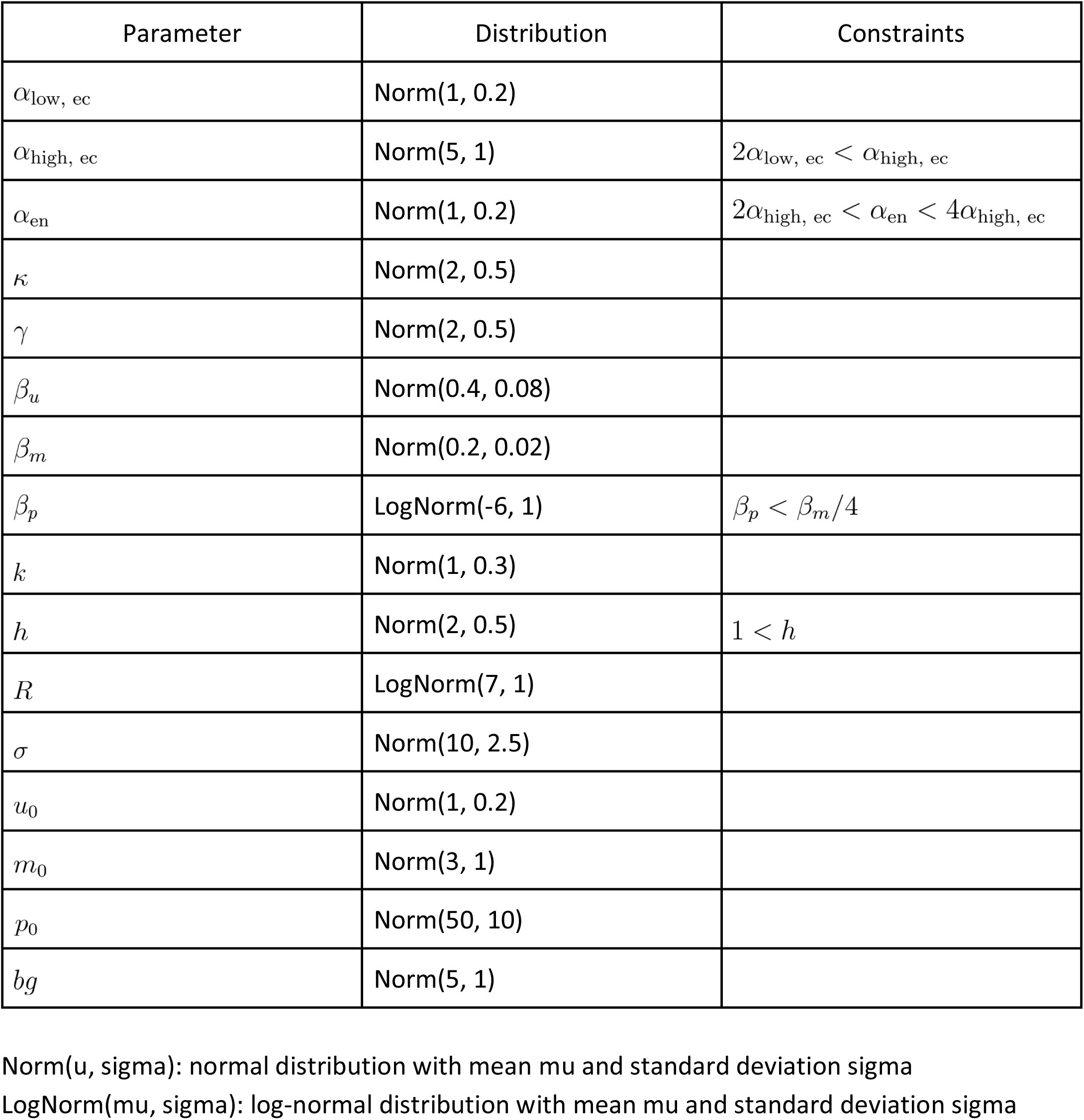

